# FocA: A deep learning tool for reliable, near-real-time imaging focus analysis in automated cell assay pipelines

**DOI:** 10.1101/2023.07.20.549929

**Authors:** Jeff Winchell, Gabriel Comolet, Geoff Buckley-Herd, Dillion Hutson, Neeloy Bose, Daniel Paull, Bianca Migliori

## Abstract

The increasing use of automation in cellular assays and cell culture presents significant opportunities to enhance the scale and throughput of imaging assays, but to do so, reliable data quality and consistency are critical. Realizing the full potential of automation will thus require the design of robust analysis pipelines that span the entire workflow in question. Here we present FocA, a deep learning tool that, in near real-time, identifies in-focus and out-of-focus images generated on a fully automated cell biology research platform, the NYSCF Global Stem Cell Array®. The tool is trained on small patches of downsampled images to maximize computational efficiency without compromising accuracy, and optimized to make sure no sub-quality images are stored and used in downstream analyses. The tool automatically generates balanced and maximally diverse training sets to avoid bias. The resulting model correctly identifies 100% of out-of-focus and 98% of in-focus images in under 4 seconds per 96-well plate, and achieves this result even in heavily downsampled data (∼30 times smaller than native resolution). Integrating the tool into automated workflows minimizes the need for human verification as well as the collection and usage of low-quality data. FocA thus offers a solution to ensure reliable image data hygiene and improve the efficiency of automated imaging workflows using minimal computational resources.

## 1. Introduction

The implementation of automated image acquisition in a laboratory setting can yield numerous advantages (Shariff et al., 2010). It helps to maintain consistency in the imaging process by eliminating the need for manual adjustments and measurements (Mallard et al., 2013), which could otherwise lead to inconsistencies and inaccuracies. When monitoring long term cell cultures, automated imaging can be useful to detect changes over time both for research and production purposes (Doulgkeroglou et al., 2020; Fischbacher et al., 2021). Furthermore, automation enables the acquisition of a larger number of images in a shorter time frame, which can increase the efficiency and speed of the laboratory’s research and production, and also enables the monitoring of a large number of samples at once (Bray et al., 2004; Fischbacher et al., 2021). This can be beneficial for large-scale cell production, such as in the case of iPSC reprogramming and antibody development (Li et al., 2010; Paull et al., 2015). Despite these enormous benefits, one downside to automated imaging is the need for real-time feedback; if quality control issues, such as exposure or focal issues, are present, data can more easily go unnoticed until it is too late, requiring layers of human supervision through the process.

Therefore, it is essential to ensure that the images acquired with automation are in focus in a timely manner. This is crucial to guarantee that the subsequent analysis is accurate, and also to provide the opportunity to re-acquire the image at the desired time-point if the image is not in focus. For this reason, many algorithms have been created to detect out-of-focus images.

Several methods for focus evaluation in cell culture images are available including the inverse coefficient of variation (ICV), normalized intensity variance (NIV), and *Power Log-Log Slope* (PLLS) (Bray et al., 2012). The ICV is the ratio of average image intensity to standard deviation of image intensity and is a unitless measure that is useful in comparing the variability of different datasets. A high ICV indicates that the mean is much larger than the standard deviation, suggesting that the dataset is less variable. Thus, a higher focal blur leads to an increase of this parameter and a higher difference in cell counts compared to the ground truth. NIV is the ratio of intensity variance to the squared average image intensity. The NIV is a dimensionless quantity that ranges between 0 and 1, with a value of 0 indicating that all pixels in the image have the same intensity, while a value of 1 indicates that the pixel intensities are highly variable across the image. An out-of-focus image, with higher blur, thus results in a smaller value. While both of these metrics offer useful approaches to assessing image quality, neither is reliable in an automated setting, as many times there are dust specks or dead cell populations floating at a different focal plane; these “artifacts” may appear very bright and thus throw off the standard deviation value and, consequently, the NIV and ICV values. Furthermore, there are many cases where the contrast is not very enhanced, but still the images are in focus. In those instances these metrics would show values that would place the images into the out-of-focus class. PLLS is the slope of the power spectral density of the pixel intensities on a log-log scale. It estimates the rate at which the amplitude of image features decreases with increasing spatial frequency, measuring effectively how defined the structures are within the image. As high-frequency image components are lost as an image is blurred, the PLLS value will thus decrease as the focal blur increases. While PLLS is less prone to error due to out-of-focus, bright objects, still the separation of in-focus from out-of-focus images in a dataset using the PLLS metric requires a user-selected threshold, making it difficult to interpret the absolute value of the metric on any given image. Identifying the absolute focus quality of a single image without any user-defined or dataset-specific threshold has been a challenging problem that remains unsolved. In fact, the need for a threshold, which may vary for each image channel, makes it impossible to perform online automated focus quality analysis while acquiring images.

To overcome these limitations, many Machine Learning (ML) algorithms have been widely used with great success (Sampson et al., 2021; Weilong Hou et al., 2015; Yang et al., 2018). The capacity of deep neural networks to learn nuanced features and outperform non-ML methods typically comes at the cost of a large initial training set and more computational overhead costs. The robust performance of classical metrics like PLLS indicates that the features contributing to focus level classification are simple and, in turn, can be learned using fewer convolutional layers than those needed for more complex classification problems. Architectures with fewer layers are also more suited to time-sensitive applications, like imaging assays in automated cell-production pipelines.

Yang et al introduced a deep learning-based strategy with a network trained on cell nuclei images that were artificially defocused into 11 focus levels, achieving an accuracy close to 95% within one focus level and comparable performance in a simpler binary in/out-of-focus classification task. The tool was made available as a free software library and as plugins for Fiji (ImageJ) and CellProfiler. While it has proven to be accurate and generalizable, applications in highly automated settings remain a challenge.

While the model’s accuracy and generalizability have been demonstrated, its application in highly automated settings remains a challenge. Nonetheless, there are specific aspects that were not well-suited for our particular case. Notably, the model’s lack of training on real data restricts its adaptability to real-world datasets, which often entail unforeseen variations.

Moreover, the extensive eleven-level categorization of blurriness, as outlined in the original paper, proves excessive for our intended purpose as a binary classifier. As we strive for seamless scalability and automation, implementing the Fiji plugin poses difficulties, primarily due to its requirement for manual input during deployment.

Here we present FocA (Focal Analysis), an ML tool designed for the near-real-time focus checking of cell culture plates in highly-automated pipelines. FocA is capable of reliably and efficiently identifying out of focus images from a fully automated assay giving immediate feedback, allowing for the re-imaging and analysis of affected (out of focus) assay plates.

Although designed for highly-automated laboratories, we believe that this tool will serve useful to the wider community in both automated and non-automated setups.

## 2. Results and discussion

### 2.1. Overview of the workflow

FocA was developed to assess the focus quality of live/dead assays used to monitor the viability of cells. In the automated workflow, the plates of cells, imaged at individual well level, are ingested and each well is assigned a focus score (Figure 1a); the decision to re-image a plate is made based on these results. In these assays, we estimate the cell count in each well based on the intensity and count of nuclei using software integrated into our imaging platform. However, in out-of-focus images, individual nuclei are indistinguishable from each other, leading to inaccurate counts and viability measurements. Additionally, the imager’s auto-focus can fail due to slight variations in plate orientation, visible scratches in the bottoms of wells, and other unexpected artifacts. FocA was optimized to efficiently verify the focus quality of large volumes of scans from small patches of heavily downsampled images (Figure 1b). Further, the tool was designed to maximize performance from an imbalanced dataset (Figure 1c) through automatic proportional data augmentation and undersampling. Since out-of-focus images are much less frequent than in-focus images, we implemented a strategy to create balanced, maximally varied training sets. Similarly, training runs for our deep learning network were structured to maintain variety within mini-batches (Figure 1c). To assess the focus well images, we divided the images into small patches, sorted them by standard deviation, and selected patches from the center of the standard deviation range. The focus of the selected patches is evaluated individually by our deep learning network and the results are combined to label the original well image as in-focus or out-of-focus (Figure 1d).

**Figure 1:**
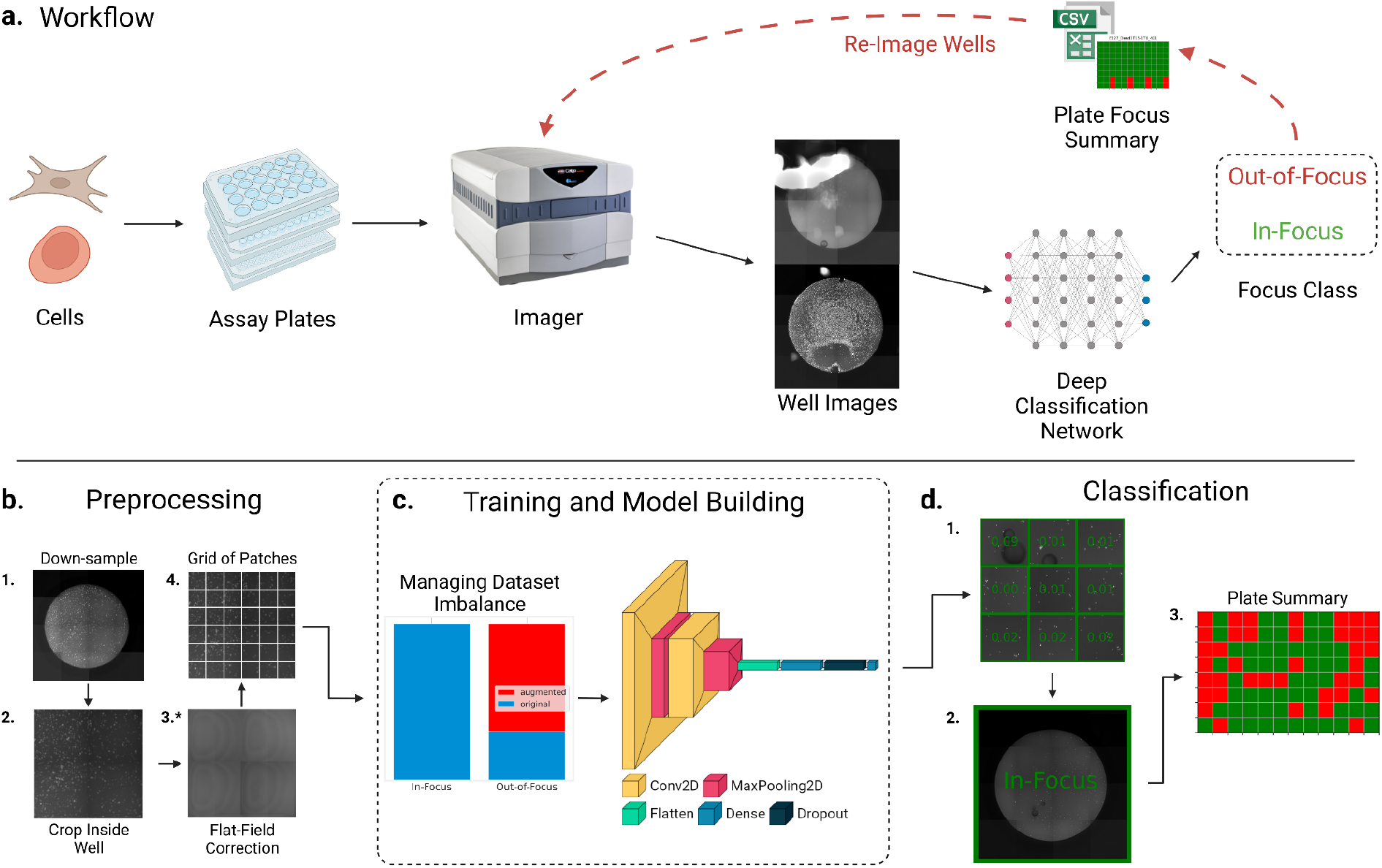
FocA is a tool that reliably detects suboptimal images in an automated workflow. **a**) *FocA’s end-to-end workflow*: Assay plates are imaged at well-level. The images are then evaluated using FocA’s deep neural network model in near-real time. The outputs are a CSV file with all the logged information about focus quality, and a visual map that shows in focus and out of focus wells. If any wells on the plate are classified as out-of-focus, the user is notified instantly to give the chance of re-imaging the plate in a timely manner **b**) *Image preprocessing*: Each well-level image is 1. exported at a downsampled resolution from the imager, 2. cropped to get read of the outer part of the well, 3. optionally flat-field corrected, then 4. split into patches to be used for training. **c**) *Building and training the deep neural network*: In order to compensate for an initially imbalance dataset, we provide a strategy for balancing and maximizing variety within the two classes. Then we built a shallow neural network to classify each patch into the binary classification in-focus or out-of-focus. **d**) *Patch classification to plate focus summary*: A subset of the patches from the well image is selected based on standard deviation to remove empty patches and other artifacts. Each selected patch is then assigned a score by the trained network and the well’s overall binary label is calculated based on a confidence-weighted mean. The well’s class is determined based on a binary threshold of 0.5. After evaluating all wells on a plate, FocA provides a visual summary to the user.

### 2.2. Training set preparation for optimal balance and variance distribution

Automatic imagers are, for the most part, consistent in their imaging ability from well to well, and plate to plate; accordingly, the majority of scans tend to be in-focus (Figure 1c). For this reason, building a balanced training set of in-focus and out-of-focus scans was challenging. The process of manually labeling each image is very time consuming, especially if a class is generally much more frequent than another. For this reason, we implemented a strategy to balance the dataset automatically and effortlessly, making it possible to overcome the issue of the class imbalance.

To do that, we used a combination of data augmentation (rotation and reflection of the patches) and undersampling of the larger class while ensuring the maximum number of plates are represented, similar to the popular SMOTE technique (Chawla et al., 2002). Using this approach, we were able to balance each class in proportion to the size of the largest class, resulting in a balanced dataset for training. We first take non-overlapping patches covering the complete area of the cropped well in a grid, then randomize the location and orientation for additional patches. For example, using the strategy above, if 36 patches per well were obtained from in-focus wells, and, in total, there are half as many out-of-focus wells, then 64 patches per well would be obtained from out-of-focus wells. In addition to this, we maximized variety within training batches by loading into memory diverse subsets of well images from different plates, which share the same acquisition settings and experimental pipeline, therefore have similar variation, and allocating a patch from each well into individual training mini-batches. These techniques allow for an effective shuffling of the training set and uniformly diverse mini-batches that minimize the possibility of over-classification.

### 2.3. Design of the convolutional neural network for speed and performance optimization

To optimize for size and speed, we experimented with different image export resolutions and patch sizes taken from well-level images. We found that larger patch sizes generally correlated with higher patch-level classification accuracy, as shown in Figure 2a, as they capture a greater percentage of the well area, leading to more cells and features contributing to the overall classification. Conversely, small patches may contain only sparse cell populations or even be empty, resulting in a lack of well variation. Lowering the export resolution, while sacrificing finer details, increased the patch-level classification accuracy as it captured a larger area for a constant patch size, leading to a better performance for smaller patch size (Figure 2a). The final configuration was chosen by evaluating the class-specific accuracies of the model when varying downsampling and patch size (Figure 2a; Supplementary Table 1). As our priority is to avoid missing any low-quality data that might invalidate subsequent automated analysis, we looked at in-focus and out-of-focus accuracies separately, and picked a configuration that would ensure the lowest false negative classifications.

**Figure 2:**
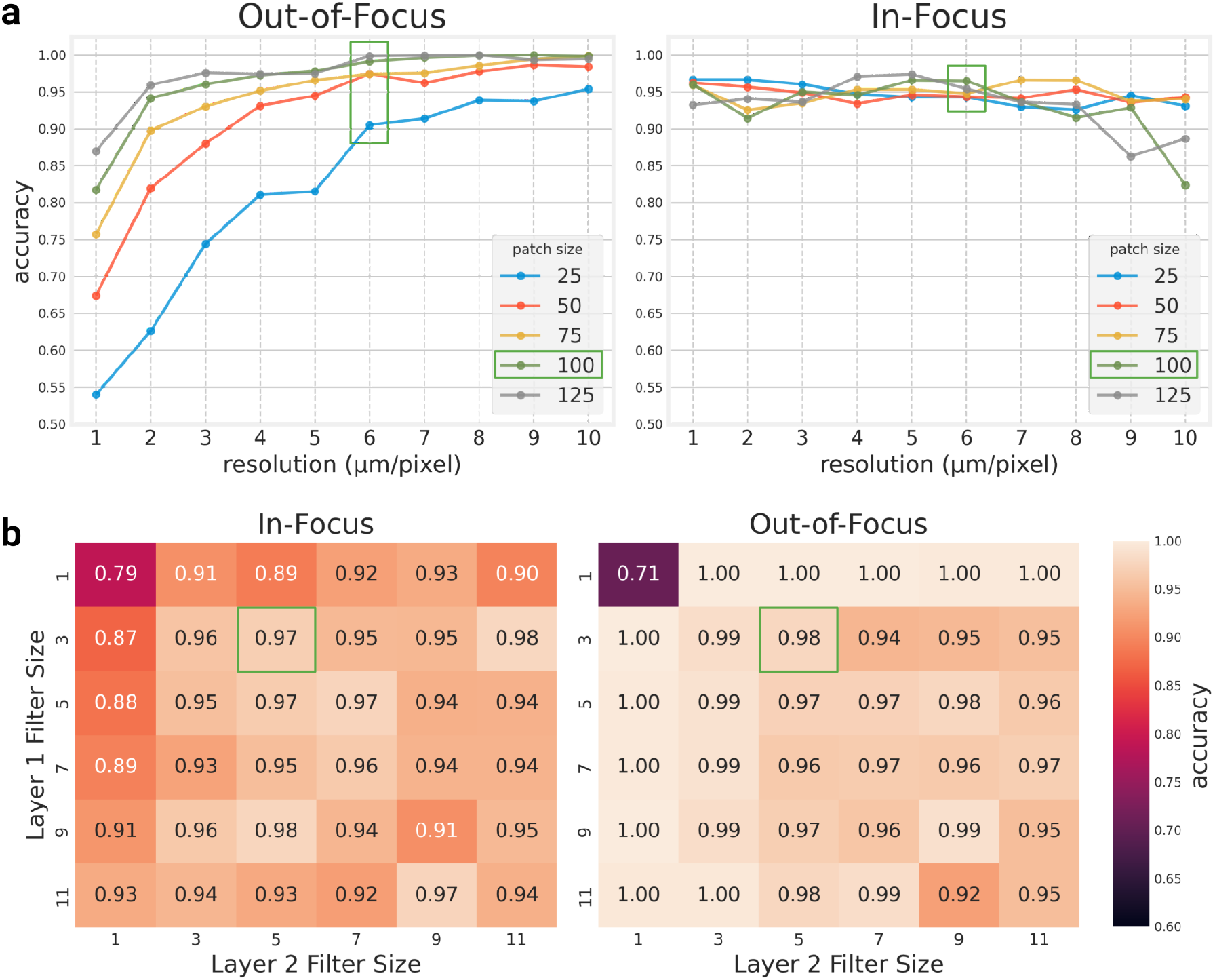
FocA’s architecture and data pre-processing is optimized for efficiency. Overview of in-focus and out-of-focus accuracy for different model and data configurations for model optimization. All results are averaged over 5 cross-validation folds. **a**) *Patch size and image resolution*: An experiment varying the size of patches extracted from images and the resolution of well images to optimize computational resources and memory usage while preserving accuracy. The final chosen configuration of 100×100 pixel patches and a resolution of 6μm/pixel is highlighted with green boxes. **b**) *Convolutional filter size accuracy heatmap:* an experiment varying the size of the convolutional filters in the two convolutional layers of FocA’s deep neural network. The final chosen configuration of 3×3 and 5×5 filters for layers 1 and 2, respectively, is highlighted with green boxes.

Besides accuracy, we wanted to ensure that the choice of downsampling and patch size would ensure reliable results, and most importantly, fast analysis in production environments. For this reason the optimization was focused not only in maximizing the accuracy, but also reducing the whole process time, by minimizing the size of the exported images and the patch size for training and evaluation, especially in the conditions where faster computing is not an option (eg. GPU availability). Taking all these factors into account, we selected to use a final resolution of 6μm/pixel resolution, and trained the dataset on patches of size 100×100 pixels, obtaining a patch accuracy of 0.986, and a computational speed of 2 seconds using a medium sized GPU, and of 3 seconds using CPUs on an entire plate (96 images) (Supplementary Figure 1a).

This initial optimization was performed on a network similar to those previously described (Yang et al., 2018). From that initial network, we compared the effects of different filter sizes for the two convolutional layers to find a configuration with optimal stability and patch-level classification accuracy (Figure 2b). The size of these convolutional filters, combined with max-pooling, determines the spatial resolution of the extracted features used in final classification of image patches. After selecting filter sizes of (3,3) and (5,5), respectively, for the two convolutional layers, we optimized the number of filters per layer, preserving a 1:2 ratio of filters between the first and second convolutional layers. We built a matrix with the accuracy of the model varying the filter size of the network both for overall accuracy and for out-of-focus images specifically (Figure 2b). To prevent overfitting and premature convergence, we employed L2 regularization on both layers and utilized a large batch with balanced classes and evenly distributed wells taken from different plates. For all these optimization tests, we used a small learning rate of 3e-5 and employed 5-fold cross-validation with 120 wells per class for training and 30 wells per class for validation. The training and validation sets were constructed from a small pool of manually labeled wells described in Section 5.5. The final network has 32 and 64 filters (Supplementary Figure 1b) in the two convolutional layers, respectively, and achieves 0.974 (0.991 for out-of-focus, 0.965 for in-focus) mean accuracy (averaged over the 5 folds) for patch level classification. The detailed diagram of the neural network can be found in Supplementary Figure 1c. The parameters for the experiments on patch size and downsampling, and convolutional filter size are available in Supplementary Table 1.

### 2.4. Well-level focus evaluation

The primary aim of FocA is to spot flaws in automated image acquisition steps without human supervision. For this reason, our aim was to minimize the chance of missing an out-of-focus well, ultimately at the cost of having more misclassified in-focus wells. Cell culture typically expresses high phenotypic differences both for how the experiment was designed, and because of natural variation. Images of very dense cells appear very different from sparse populations. Furthermore, there is often debris or floating objects (e.g. dead cells) that can cause images to appear out-of-focus when viewed out-of-context. For this reason simply taking a random or sequential subset of patches from the well can be misleading. To tackle these observations, we partitioned the well-level image into a grid of separate patches and ranked them based on the standard deviations of their pixel intensities. This enabled us to give priority to sections of the cropped well-level image that were not empty, as well as avoid sections that contained bright artifacts that could affect the classification (Figure 3a). We next used our model to evaluate the focus of individual patches with the standard deviation closer to the middle of the range, before assigning a well focus classification. To evaluate this approach, we assessed the accuracy based on a held-out set of 60 wells (30 each in-focus and out-of-focus). Accuracy was calculated as the mean binary prediction accuracy of 16 patches taken from each of these 60 wells.

**Figure 3:**
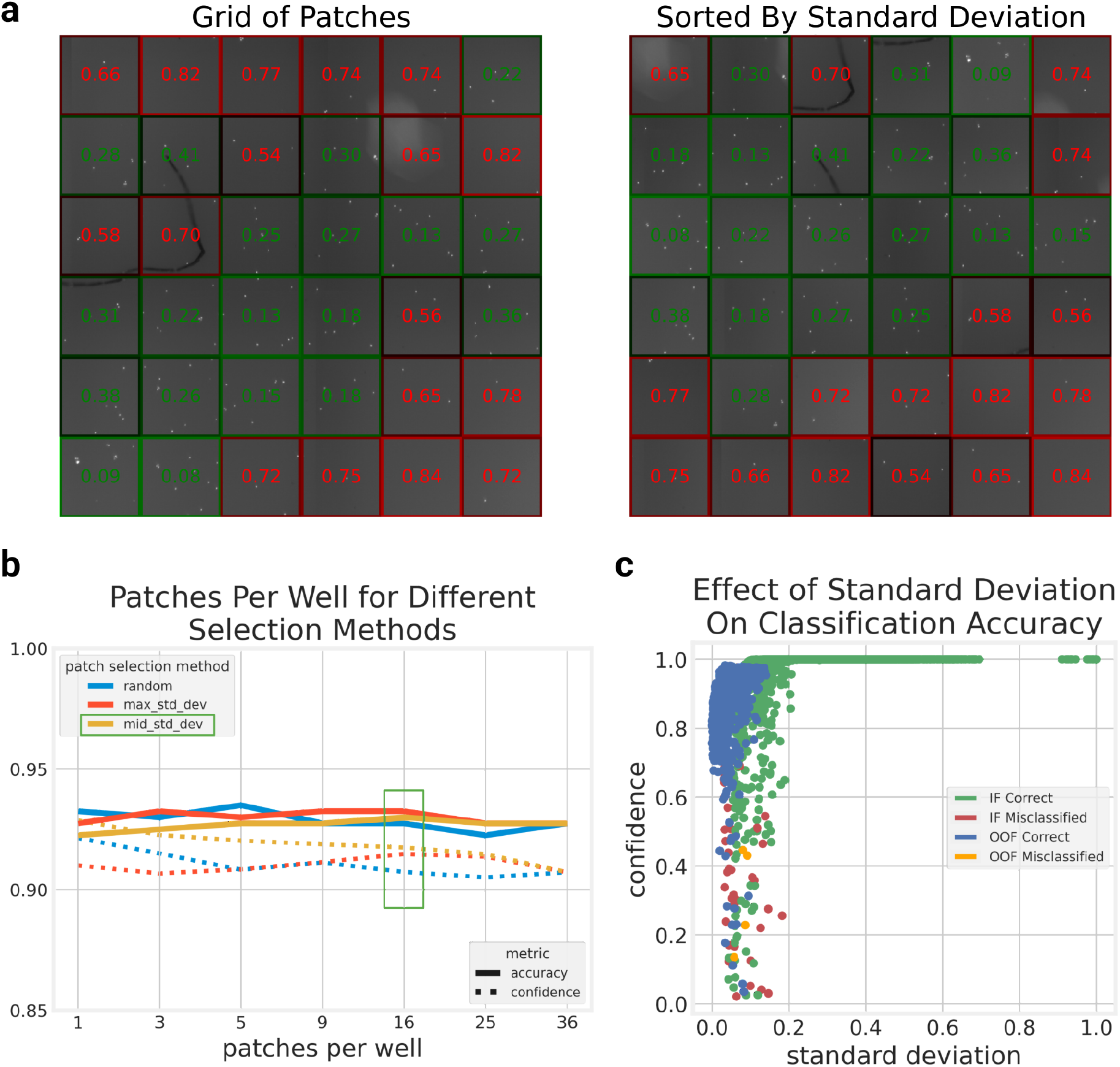
FocA classifies wells as in-focus or out-of-focus with excellent accuracy and high confidence. **a**) *Sorting patches by standard deviation*: Visualization of scores assigned to patches in their original organization and sorted in descending order of standard deviation. Generally, patches containing artifacts and anomalies will be placed at the top of the standard deviation range and empty or sparsely populated patches will be placed near the end of the range. The borders of the patches are colored according to confidence and class, where green is in-focus, red is out-of-focus, and brighter borders are more confident. **b**) *Patch selection methods and well classification confidence*: An experiment testing methods for selecting patches from a complete grid and the number of patches required for confident well classification. The final selection of 16 patches per well and the “mid_std_dev” method–taking patches from the middle of the sorted standard deviation range–is highlighted with a green box. The “max_std_dev” method indicates patches with the highest standard deviation were selected first and the “random” method indicates patches were randomly selected from the range. Results are computed over 400 held-out wells (200 in-focus, 200 out-of-focus). **c**) *Patch standard deviation and classification confidence*: A visualization of how patch classification confidence is impacted by standard deviation. Each point is a single patch and the plot contains 400 held-out wells (200 in-focus, 200 out-of-focus) with 16 patches taken for each well using the “mid_std_dev” technique mentioned in b). For this plot, the standard deviations have been normalized to the range [0,1].

Each patch was assigned a single value in the range [0, 1], where 0 is in-focus and 1 out-of-focus. Using the selected 100×100 pixel patch size, we divided the well into 36 non-overlapping patches. Figure 3a shows an example of a well with overlaid prediction score for each patch, and while this well is in focus it presents some empty patches and some high intensity objects in a small subset. Using the entire subset of patches and/or selecting a random subset would have potentially led to a misclassification. For this reason, and to minimize the amount of data to be analyzed, we evaluated how many patches were necessary to achieve the best prediction.

To classify the well image we made sure that the patches with prediction scores closer to a random chance were weighted less than the ones with higher confidence (De Luca et al., 2023). We opted to use a metric that takes into account the uncertainty and confidence of the model’s predictions, rather than simply averaging the values to ensure that the automated pipeline would not overlook any out-of-focus wells by assigning higher importance to patches where the model exhibited greater certainty. By doing so, we aimed to minimize the risk of the misclassifications that occur in response to fuzzier patches, which are typical in cell culture images.

We implemented a strategy using the confidence metric, measured as the prediction score value’s absolute distance from the chance value 0.5; we assessed the well-level image focus by taking the confidence-weighted average of the patch predictions before classifying the well as either in-focus or out-of-focus based on a threshold of 0.5. We then looked at the performance for randomly selected, locally ordered, and standard deviation sorted patches, varying the number of patches in each group (Figure 3b). We found the optimal number of patches to be 16 taken from the middle range of standard deviation, although any number of patches between 9 and 36 provided similar results (Figure 3b).

We next performed a further evaluation and looked at the confidence of the prediction of each patch in relation to the standard deviation. As expected, the higher the standard deviation, the higher the confidence that the image is in focus. We can see this result both at patch level (Figure 3c) and well level (Supplementary Figure 3). Representative examples of common types of wells and their prediction scores at patch level are shown in Supplementary Figure 2. Exact experimental configurations used to produce results in Figure 3 are listed in Supplemental Table 1. All results detailed in this section, Figure 3, and Supplementary Figure 2 and 3 were generated from a model trained on a larger pool of images selected deploying FocA in our automated pipelines. Specific details are included in Section 5.5.

### 2.6. Generalizability: testing the same framework across different datasets

To ensure the generalizability of FocA’s framework and selected parameters across different use cases and image types, we generated a dataset comprised of plated fibroblasts stained with two different dyes (a nuclear stain and a membrane stain) and captured at two different magnifications (10X and 20X). Our microscope allowed us to image the same planes in each well, enabling the production of one in-focus image and two levels of out-of-focus images for the same field of view.

In our first experiment, we trained FocA on an increasing number of wells and evaluated its performance on a held-out validation set. Each well contained one field, which we divided into 100 patches. For the validation set, we used 32 patches per well, selecting the middle distribution of standard deviation as mentioned earlier. Since the data in this case was imbalanced towards the out-of-focus class, we employed the balancing strategy outlined in a previous session.

The results, depicted in Figure 4a, demonstrate that only a small number of images are needed to achieve good accuracy at the patch level. The optimal number of images for the highest accuracy was around 100, reaching approximately 90%. The reason for the plateau in accuracy is that, due to the images’ higher level of detail compared to the Celigo images, both in the in-focus and out-of-focus cases, some patches within the in-focus images can still appear blurry but are labeled as in focus. An example field with corresponding patch predictions is shown in Figure 4b. However, the majority of patches within the wells usually belong to the correct class, leading to correct predictions at the image level. Note that in this experiment though we kept the patch size that was optimized on images with much lower magnification to demonstrate generalizability. Optimizing the patch size on images with very different magnification would most probably increase the accuracy significantly.

**Figure 4:**
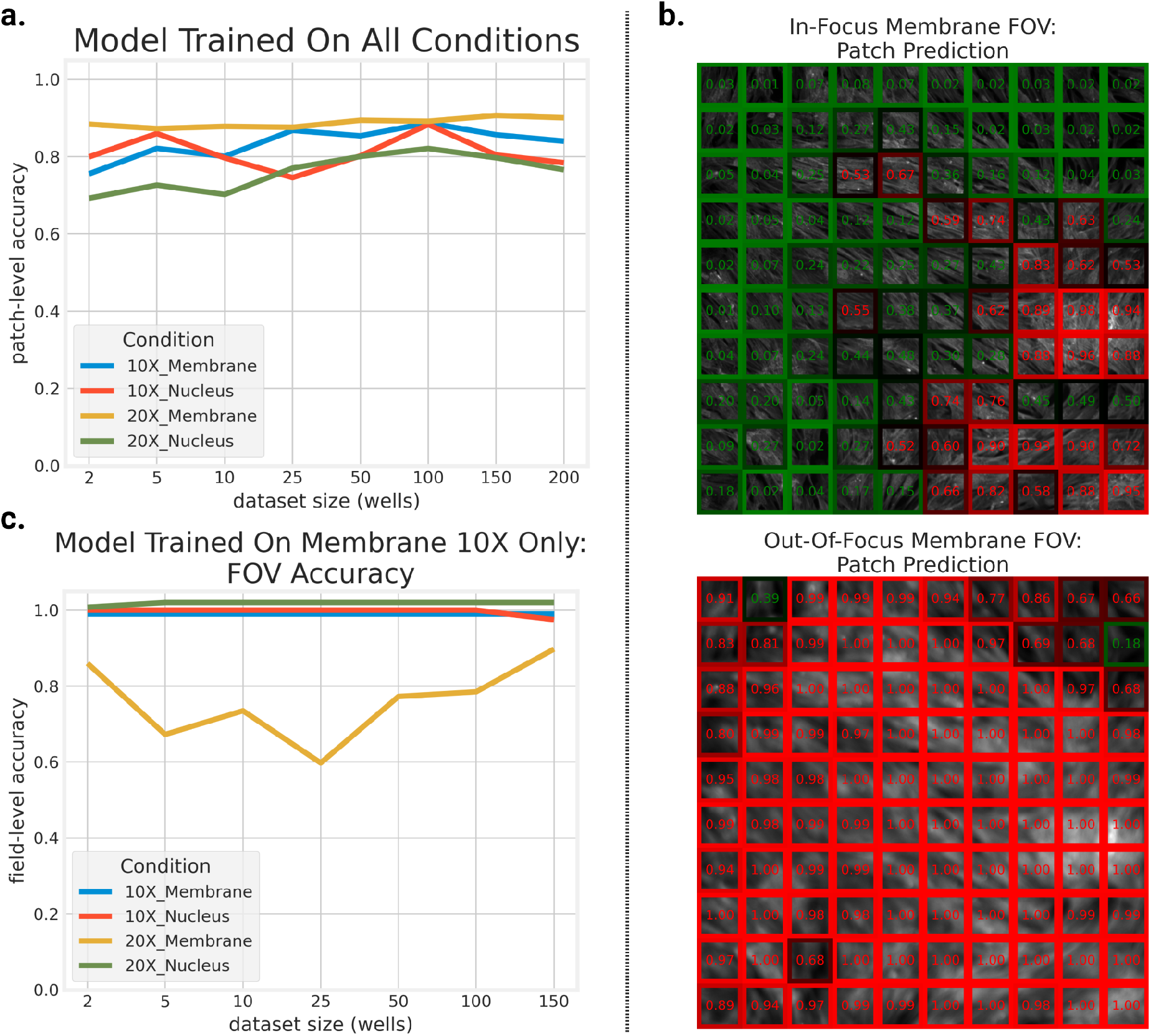
FocA generalizes to a variety of imaging configurations. **a**) *FocA can assess focus for a variety of magnifications and stains with small amounts of data:* After training on progressively larger datasets of two different magnifications and stains, FocA is proven capable of analyzing multiple stains and magnifications with high accuracy. **b**) *Patch-level classification of cells stained with WGA and Phalloidin:* Using the model from Fig 4a, we generate patch-level visualizations of one in-focus (*top*) and one out-of-focus (*bottom)* 10X image of cells stained with WGA and Phalloidin. The patches are assembled into a 10×10 grid of patches, with borders corresponding to classification score and confidence. Green and red correspond to in-focus and out-of-focus classification, respectively, while dark and bright borders correspond to low and high confidence. **c**) *FocA generalizes to new magnifications and stains:* After FocA is trained on only images of cells stained with WGA and Phalloidin at 10X magnification, it achieved high accuracy in classifying unseen images captured with same parameters but also images of cells stained with Hoechst at two different magnifications. Classification accuracy for individual stain/magnification combinations is presented at field-level.

Next, we investigated whether training on one condition could yield accurate predictions on other conditions. We selected the membrane channel at 10X magnification, increased the number of wells in the training set, and evaluated performance on the same validation set used in Figure 4a. Figure 4c presents the model’s performance at the field level. Once again, the predictions rapidly reached high accuracy for all dyes except one. The slightly lower accuracy for the membrane channel at 20X magnification is attributed to the highly zoomed-in nature of the images, which introduces more fuzziness in the patches compared to the smaller magnification and nuclear images, which are more compact and well-defined.

Through this experiment, we not only demonstrate that FocA can be applied to datasets with significantly different appearances compared to our automated pipeline’s output, but also that there is a considerable level of generalizability when training on one experiment type and testing on another. As a final confirmation of this, we utilized the same network described in Figure 4c on the validation set from the Celigo images, achieving a well-level accuracy of 0.78.

The results of this experiment provide strong evidence of FocA’s ability to adapt to diverse datasets and exhibit generalizability when trained on one experiment type and tested on another.

### 2.7. Deployment in an automated laboratory setting

FocA was developed to fill a real need in an automated laboratory setting. For this reason after building an optimized and reliable model, we needed to integrate it into our pipeline and make sure it would work as needed without human intervention. To integrate the tool into our cell production pipeline, we automated the analysis in a workflow shown in Figure 1a. The scans are exported from the Celigo imager to a local server where a process is executed to run a background bash script scheduler of type Cron on a two minutes basis. If new images are detected, the script initializes the FocA model to process them. The outputs of the pipeline are PNG image showing out-of-focus wells, which is sent instantly to the scientists via a messaging system, and a CSV summary of all data collected (example shown in Supplementary Table 2) from the analysis, including in-focus/out-of-focus counts, labels, and metadata such as the time of scan and analysis, and the plate name. This file is stored in an internal database. The speed of execution and subsequent text alert are crucial, since cells are live entities and can change extensively in a limited amount of time, making it hard to compare specific timepoints if the plates are not readily rescanned.

## 3. Conclusions

FocA is an imaging focus tool that concentrates on speed, reliability and ease of use; its performance enables it to be deployed routinely in an automated setting, without supervision. While other tools focus on detecting the differences in blur between images, or the levels that can occur within a dataset, our tool uses a simple quality threshold that facilitates the detection of suitable and unsuitable images for downstream analyses. FocA’s downsampling and patch-based model makes it easy to deploy and scale, and its simple code structure allows easy retraining, aiding in repurposing the same architecture for other image-quality tasks. FocA’s architecture also has the potential to create models for a variety of image quality assays, including over-exposure, empty well detection, and contamination detection. FocA’s flexibility, reliability and limited size makes it a great candidate for multiple applications that can facilitate automated workflows in cell culture laboratories of various sizes.

## 5. Materials and methods

### 5.1. Images collection

Images were acquired using the Nexcelom Celigo Image Cytometer at 4× magnification, an LED-based fluorescent blue channel used for DAPI with an excitation of 377/50, a dichroic filter of 409, and an emission of 470/22. The native resolution of the images is 7500 x 7500 pixels, 1 micron per pixel at 8-bit depth per pixel in grayscale. The images were manually sorted into two classes: in-focus (IF) and out-of-focus (OOF) exported at 1 micron per pixel resolution for the initial model building part, and at 6 microns per pixel for deployment. The Celigo images 16 fields for each 96-well-plate well then stitches these images in a 4×4 grid to form the final well image. In this paper, we used images of cells stained with Hoechst (ThermoFisher, H3570) in 96-well plates.

### 5.2. Preprocessing

The flat-field image is calculated by taking the mean projection of well-level images from an empty plate. Images were cropped to half the original width and height around the center pixel, (inside the circular well border) to remove irrelevant information. Normalization was performed by taking the minimum and maximum intensity values across all cropped images in our dataset, before normalizing over this range ([6, 255] for our 8-bit images).

Images used in the generalizability experiments detailed in Figure 4 were pre-processed differently than images collected for other experiments. Intensity values of DAPI images were truncated above the 99th percentile to mitigate noise contributions. Field images were normalized individually using minimum-maximum normalization, converting them from a 16-bit depth, to the range [0,1]. In contrast to images described above, only one field was acquired for each well.

### 5.3. Deep learning network and computational resources

The tool was deployed on a desktop computer running Ubuntu 22.04 LTS equipped with 32 GB of RAM, an i7-10700 8-core Intel CPU, and an Nvidia GeForce RTX 2080 Ti GPU. The final model configuration consists of a convolutional layer with 32 filters of size 3 x 3, a 2 x 2 max-pooling layer, a convolutional layer with 64 filters of size 5 x 5, a 2 x 2 max-pooling layer, a fully-connected layer with 512 units, a dropout layer with probability 0.4, and a fully-connected layer with 1 unit. The model was implemented in Tensorflow v2.9.0 (Abadi et al., 2016).

### 5.5. Model Training Strategy

Our dataset is split into two classes named in-focus (IF) and out-of-focus (OOF). For each of the experiments outlined in Figure 2, we trained models using an 80:20 training-validation split over a maximum of 30 epochs, using early stopping with a patience of 3 epochs, a minimum validation loss delta of 0.01, and a learning rate scheduler that decreased the learning rate by 10% each epoch. Each model used binary cross-entropy as its loss function. We trained models of the same configuration over 5 cross-validation folds. We used a batch size of 32 patches (for a total of ∼50MBs) and took 36 patches from each well image. Our initial image parameters consisted of 5μm/pixel resolution images, 36 100×100 pixel patches from each image (16 patches for validation and testing). For the initial model parameters, we fixed input size to 100×100, and the number of convolutional filters to be 32 and 64, respectively.

Our final training was conducted in 336 batches of 32 patches on a dataset of 150 each of in-focus and out-of-focus wells; this model was used to generate results in Figure 3 and Supplemental Figures 2 and 3. To clarify the changes in parameters between experiments we provide Supplementary Table 1, documenting the entire configuration for all experiments, with experiment columns ordered chronologically. The first four columns remain mostly consistent aside from the tested parameters which are marked *Varied*. Experiments in columns 1 through 3–*Convolutional Filter Size*, *Number of Convolutional Filters*, and *Patch Size vs Image Resolutions*–each depend on the previous experiment’s results and subsequent parameter changes reflect this. For the remaining experiments, we used the optimized configuration derived from the first 3 experiments (columns 1-3), but trained on our entire dataset without a validation set using a learning rate identified from a previous experiment for a single epoch. For the *Patch Selection Method* and *Confidence vs Standard Deviation* experiments, we tested on a set of 200 random held-out wells we gathered while using the tool in our automated pipelines. Results from the *Confidence vs Standard Deviation* experiment are visualized at patch-level and well-level in Figure 3c and Supplemental Figure 3, respectively. For the last experiment on *96-Well Plate Computation Time*, we used a random set of 960 wells (10 plates worth of wells).

Our data was split into a small initial set and a larger set accumulated while FocA was deployed in our automated pipelines. This small initial set was constructed from a pool of 47 unique plate scans, 27 in-focus and 16 out-of-focus, with 1168 and 367 wells, respectively. Plate scan variation within each class was maximized using the strategy outlined in section 2.2. The testing set is kept consistent across experiments and is composed of 60 wells, 30 for each class, randomly selected from the larger set of 269 plate scans, 190 in-focus and 79 out-of-focus with 9,807 and 459 wells, respectively. Optimization experiments detailed in Figure 2 and Supplemental Figure 2 were trained and validated on the small initial set and were tested on the test set described above. For experiments detailed in Figure 3 and Supplementary Figures 2 and 3, models were trained on a subset of 150 wells from the larger set and tested on 200 held-out, randomly selected wells from the same set.

A separate dataset captured using a different imaging configuration, described in Section 5.9, was used to test generalizability. The dataset consisted of 230 wells imaged at 10X and 20X magnification for a single field-of-view, each at three different focal planes (1 in-focus, 2 out-of-focus), and 2 channels (nucleus, membrane) for a total of 2,370 images. For Figures 4a-b, models were trained on progressively larger subsets of wells imaged at all combinations of channel and magnification configurations. 100 100×100 patches were taken from each well in each of the 2 out-of-focus planes and 200 patches were taken from the in-focus plane to ensure balance across the two classes. For simplicity, we treat images captured at 2 different magnifications as two different plates for a total of 460 unique wells. When training images were added iteratively so that images from the previous set were included in the next (e.g., images from the set used to train at 100 well images, were also included in the set of 150 well images). Images from 40 randomly selected wells from the two plates (magnifications) each were held-out for validation and testing. For Figure 4c, models were trained on progressively larger sets of wells using the same method described above except that images were only taken from a single configuration: membrane channel at 10X magnification. Images from 40 randomly selected wells each were held-out for validation and testing. For testing and validation sets, 32 100×100 patches selected from the center of the standard deviation range were taken from each image. We increased the number of patches from 16 to 32 because this new dataset consisted of higher resolution images (648 x 648 vs 1080 x 1080 pixels). For Figure 4a and 4c, testing accuracy for individual configurations is plotted separately in different colors. The results pictured in 4b are generated from the model trained on 100 wells from Figure 4a.

For the generalizability experiments detailed in Figure 4, all models were trained using the same parameters, with the exception of batch size. Batch size was scaled linearly with dataset size, which ranged from 6 to 256 patches for models generated for Figure 4a and from 6 to 32 patches for Figure 4c. Training for each model was performed for a maximum of 100 epochs, with an early stopping minimum validation loss delta of 0.01 and a patience of 10 epochs. Aside from an increase in epochs and patience and scaling the batch size with dataset size, the training hyperparameters remained the same as those used to train models for the experiments detailed in Figures 2 and 3.

### 5.6 Patch to Well Classification

To determine the focus of well images, we use a technique that uses the confidence-weighted average of selected patches. Specifically, we calculate the confidence value of each patch’s classification score, multiply each score by its corresponding confidence value, sum the products, and divide by the sum total of all the confidence values. We then threshold the final weighted average by 0.5 to classify the image as either in-focus or out-of-focus.

### 5.7. Example deployment pipeline

Scans are exported from the Celigo imager to a Network Attached Storage (NAS) device for long term storage. Working on a centralized computer, running a Unix operating system, we utilized a background job scheduler (Crontab) to execute the StartFocA.sh script on a routine interval. The script initializes an Anaconda environment and runs a Python script. In the case all the scans in our NAS have already been analyzed, the process closes without generating any outputs, nor initializing the main FocA script. If new scans are detected the script will process them and produce two files: a png image of a visualization of each of the analyzed plates, and a csv file summarizing all the data we collected through the analysis(count of OOF and IF wells, time of scan, plate name). Application specific APIs are used to provide real-time feedback to scientists using cloud-based messaging platforms, as well as feed data into centralized databases through tools such as AWS Lambda.

### 5.8. Wet lab experiments detail

The NYSCF Global Stem Cell Array (NYSCF GSA) is a bespoke platform of interconnected robotics and peripherals to facilitate the automated culturing of human cells, including induced Pluripotent Stem Cells (iPSCs). The complex orchestration of Array machinery to accomplish a thaw, passage, or freeze is colloquially known as a “method.” All array methods in which cells enter suspension employ Dead/Total Plates (DTPs) to generate an approximation of the total number and viability of cells being subsequently seeded or frozen. The process by which Automated DTPs are used within the GSA to generate accurate records and perform seeding density calculations without the need for human interaction is briefly described here: Perkin Elmer ViewPlate-96 Optically Clear Bottom, Tissue Culture Treated, Sterile plates are processed with 90μl per well of a count solution consisting of, per 10 mL of CountSolution, 10, L Phenol Free DMEM media (ThermoFisher, 21041025) with 5% BSA fraction V (ThermoFisher, 15260037), 25μL Hoechst 33342 1 mg/mL (ThermoFisher, H3570), 7.4 μL Propidium Iodide 1.5 mM (ThermoFisher, P3566), and 1.11 μL Thiazovivin (THZ) at 10mM (SigmaAldrich 1226056-71-8). These DTPs are then seeded with an aliquot of cell suspension within a method, typically at a volume of 10 μL. DTPs are then transferred via a robotic arm to a Cytomat24 (ThermoFisher, 51033211) for an incubation period of 10 mins. Post incubation, DTPs undergo brief centrifugation using an Agilent VSpin at 13 g for 5 mins to ensure all cells are attached to the bottom of the plate and within the same focal plane. The DTP is then transferred via a robotic arm to a Celigo Image Cytometer for automated image acquisition before images are exported.

### 5.9. Generalizability Experiments

A 384-well imaging plate (PerkinElmer 6057300) of primary human fibroblast was generated as previously outlined (Paull et al., 2015). The plate was fixed and fluorescently labeled using an automated liquid handling system (Hamilton STAR). It was fixed with 16% Paraformaldehyde (Electron Microscopy Sciences, 15710-S) and washed with 1× HBSS (Thermo Fisher Scientific, 14025126). The 384-well plate was then permeabilized in 0.1% Triton X-100 (Sigma-Aldrich, T8787) in 1× HBSS. The Hamilton system washed the plate twice before incubating the cells in a cocktail containing Alexa Fluor® 568 Phalloidin (Invitrogen™ A12380), Hoechst 33342 trihydrochloride (H3570, ThermoFisher), trihydrate (Invitrogen™ H3570), and Molecular Probes Wheat Germ Agglutinin, Alexa Fluor 555 Conjugate (Invitrogen™ W32464) at room temperature for 30 minutes. The plate was then washed three times with 1× HBSS, sealed with 1× HBSS in the wells, and refrigerated until removal for imaging the next day.

The 384-well plate was then imaged using the Opera Phenix^®^ High Content Screening System (PerkinElmer). The inner 240 wells of each plate were imaged using the confocal objective on both the 10× (10× Air, NA 0.3) and 20× magnification (20× Water, NA 1.0). We imaged one field per well. The plate was then imaged at three Z-planes across both magnifications. Since the Opera Phenix™ has a laser-based autofocus system, the plate had to be imaged at several focal planes to artificially induce “out-of-focus” images. The plate was imaged at 6.5, 26.5, and 46.5μm heights on the z-drive at the 10× magnification to generate images of an in-focus, partially out-of-focus, and fully out-of-focus well respectively. The focal planes varied for the 20× magnification, using which the plate was imaged at –2.0, 6.5, and 15μm to respectively recreate in-focus, partially out-of-focus, and fully out-of-focus images. To capture the images of the plate, two different channels were used with differing combinations of excitation and emission spectra: 375 nm and 435-480 for Hoechst 33342, and 561 nm and 570-630 for WGA and Phalloidin. 2× binning was performed on each image.

## Code and Data Availability

All code is available under Business Source License 1.1 and shared through a GitHub repository (https://github.com/NYSCF/foca_release). The code is divided into three main directories: train, models, and deployment. The train and deployment directories hold functions and classes necessary to train and deploy the tool, while the models directory holds the implementation of FocA’s deep learning network. Functions for the creation of variation-optimized, rebalanced datasets can be found in the dataset module of the train directory. While our variance-optimization methods depend on an image naming scheme with plate and well information, we offer another method designed to work on generic datasets. Training and data parameter configurations for the experiments we ran and used for model training are housed in YAML files in the *experiment_configs* subdirectory in *train.* The user can fill in their desired parameters (defaults are also given) and train a model by passing their dataset CSV filename and the desired YAML config file. More instructions and details are included in the repository’s README. The authors provided detailed instruction plus notebook implementation for easier use both for training the model and deploying only.

## Supporting information

Supplemental Table 1 + Sample Output

## Acknowledgements

We thank members of the NYSCF Global Stem Cell Array® Team for cell handling and method development that led to the experiments. We are grateful to Sandra Capellera-Garcia, Corvis Richardson and Raeka Aiyar for manuscript input and review. All figures were created using Biorender.com.

## Figure Captions

Supplementary Table 1: Experiment Training and Data Parameters

Parameters used in data preprocessing and deep learning network training for each result presented in the figures.

Supplementary Table 2: Sample Output

An example of the tabulated statistics output by FocA for two plates.

**Supplementary Figure 1:**
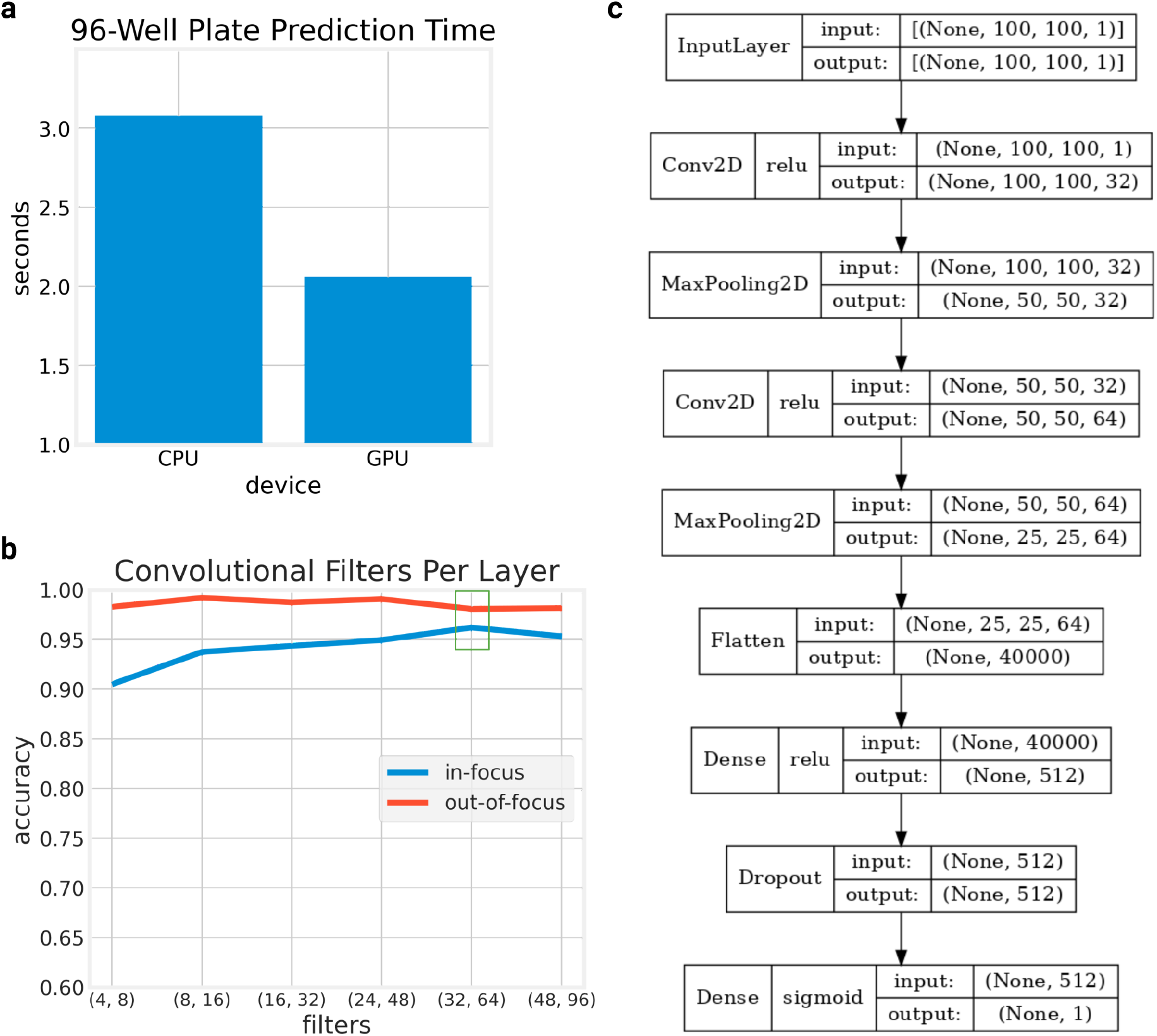
FocA classifies well images in near-real-time using either CPU and GPU processing. **a**) *Prediction time*: Time in seconds to compute predictions for 96 wells using FocA on a CPU and GPU. Results are averaged over 10 plates (960 wells). **b**) *Optimization of the number of filters per convolutional layer*: Experimental results to determine the optimal number of filters in the deep neural network’s two convolutional layers based on accuracy. The chosen configuration of 32 and 64 filters is highlighted with a green box. **c**) *Model Architecture*: Detailed diagram of the model’s architecture including layer types, activation functions, and input and output shapes.

**Supplementary Figure 2:**
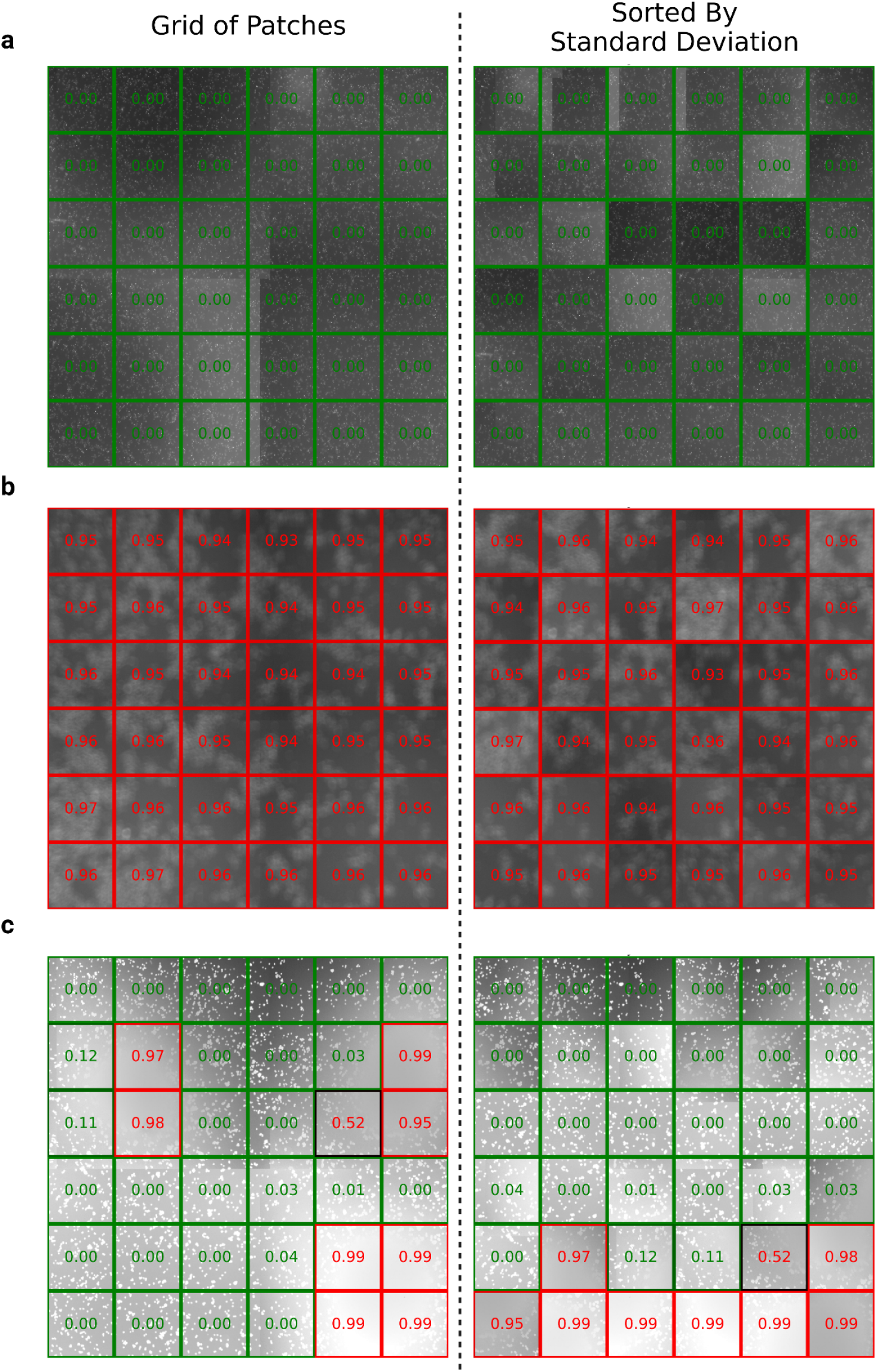
FocA classifies well images of different focus qualities accurately. Examples of three different wells with patch borders and scores overlaid. Brightness of the border corresponds to confidence of classification where darker is less confident, red is out-of-focus, green is in-focus. **a**) *In-focus well* **b**) *Out-of-focus well* **c**) *Over-exposed in-focus well*

**Supplementary Figure 3:**
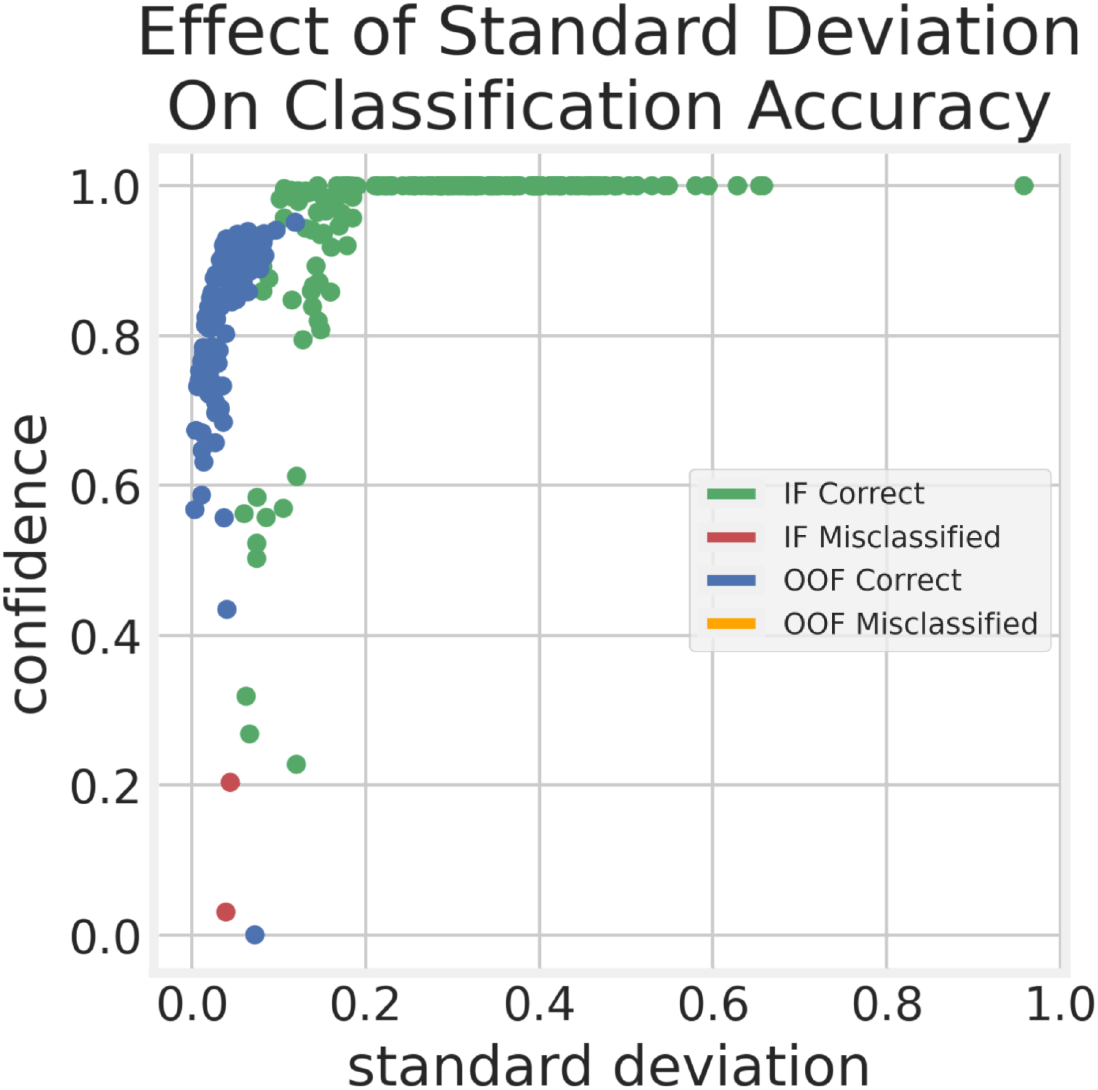
FocA is extremely unlikely to misclassify out-of-focus wells. Well-level results of the confidence vs standard deviation experiment. Alternative representation of results from Figure 3c where each point is computed by averaging the confidences and standard deviations for the patches in a well. For this plot, the standard deviations have been normalized to the range [0,1].

